# Disconnection somewhere down the line: Multivariate lesion-symptom mapping of the line bisection error

**DOI:** 10.1101/2020.04.01.020271

**Authors:** Daniel Wiesen, Hans-Otto Karnath, Christoph Sperber

## Abstract

Line Bisection is a simple task frequently used in stroke patients to diagnose disorders of spatial perception characterized by a directional bisection bias to the ipsilesional side. However, previous anatomical and behavioural findings are contradictory, and the diagnostic validity of the line bisection task has been challenged. We hereby aimed to re-analyse the anatomical basis of pathological line bisection by using multivariate lesion-symptom mapping and disconnection-symptom mapping based on support vector regression in a sample of 163 right hemispheric acute stroke patients. In line with some previous studies, we observed that pathological line bisection was related to more than a single focal lesion location. Cortical damage primarily to right parietal areas, particularly the inferior parietal lobe, including the angular gyrus, as well as damage to the right basal ganglia contributed to the pathology. In contrast to some previous studies, an involvement of frontal cortical brain areas in the line bisection task was not observed. Subcortically, damage to the right superior longitudinal fasciculus (I, II and III) and arcuate fasciculus as well as the internal capsule was associated with line bisection errors. Moreover, white matter damage of interhemispheric fibre bundles, such as the anterior commissure and posterior parts of the corpus callosum projecting into the left hemisphere, was predictive of pathological deviation in the line bisection task.

## 1. Introduction

The line bisection task is a widely used test in the diagnosis of spatial perception deficits after stroke. Originally, this task was introduced to assess visual field defects (for a review see Kerkhoff and Bucher, 2008) and later adopted in the diagnosis of spatial attention deficits (Schenkenberg et al., 1980), where it quickly became established as a routine test in neuropsychological test batteries (e.g. Halligan et al., 1991; Vaes et al., 2015). In the line bisection task, the patient is asked to manually mark the midpoint of a horizontally presented line. A deviation from the true midpoint to the ipsilesional side is typically seen as a sign of post-stroke deficits in spatial perception.

The neural correlates of line bisection errors (LBE) have been the subject of several studies that utilised either statistical lesion behaviour mapping (Kenzie et al., 2015; Molenberghs and Sale, 2011; Thiebaut de Schotten et al., 2014; Toba et al., 2017, 2018; Verdon et al., 2010) or descriptive topographical methods (Binder et al., 1992; Golay et al., 2008; Rorden et al., 2006). Most often, LBEs have been associated with damage to the posterior parietal lobe (Binder et al., 1992; Kenzie et al., 2015; Molenberghs and Sale, 2011; Rorden et al., 2006; Thiebaut de Schotten et al., 2014; Toba et al., 2018, 2017; Verdon et al., 2010).Other critical regions were found in the posterior part of the temporal lobe or the temporo-parietal junction (TPJ) (Kenzie et al., 2015; Rorden et al., 2006), frontal lobe (Thiebaut de Schotten et al., 2014), and parts of the occipital lobe (Binder et al., 1992; Kenzie et al., 2015; Rorden et al., 2006; Toba et al., 2018, 2017). Further, several studies have suggested a critical role of white matter damage (Golay et al., 2008; Thiebaut de Schotten et al., 2014; Toba et al., 2018, 2017; Verdon et al., 2010). The relevance of white matter tracts has also been highlighted by studies using either fibre tracking (Vaessen et al., 2016) or region of interest-based multivariate lesion analysis in left hemisphere stroke patients (Malherbe et al., 2018). Especially damage to the superior longitudinal fasciculus (SLF) (Malherbe et al., 2018; Thiebaut de Schotten et al., 2014, 2005; Toba et al., 2018, 2017; Vaessen et al., 2016) and the arcuate fasciculus (Malherbe et al., 2018; Thiebaut de Schotten et al., 2014) was found to underlie LBEs. Subcomponents of the SLF connect frontal areas, such as the middle frontal gyrus and pars opercularis, with parietal areas, such as the angular gyrus and the supramarginal gyrus (Thiebaut de Schotten et al., 2011a; Wang et al., 2016).

In conclusion, while there is considerable correspondence between findings in previous studies, it is not yet possible to find a unifying theory. A likely explanation for the different findings is that rather than damage to a single anatomical module, damage to a network underlies LBEs. The relevance of brain connectivity for most cognitive functions is well known (e.g. Godefroy et al., 1998; Catani & ffytche, 2005) and has also been postulated to be relevant for spatial attention (Bartolomeo, 2006; Bartolomeo et al., 2007; Karnath, 2009; Karnath& Rorden, 2012). However, two exceptions aside (Malherbe et al., 2018; Toba et al., 2017), previous investigations used univariate topographical mapping approaches to investigate the neural correlates of line bisection. These come with methodological caveats in the identification of complex neural correlates of behavioural functions, especially with respect to brain networks (see Sperber, 2020; Sperber et al., 2019b). The two multivariate topographical studies both used an analysis approach based on game theory and included only a few brain regions of interest at once. Such a priori feature reduction by testing only a few possibly crucial hubs can be necessary due to computational or statistical limitations. However, the selected parcellation can differ from the relevant functional parcellation of the brain as well as the typical anatomy of stroke lesions. In contrast, voxel-wise analysis approaches are able to maximize an analysis’ ability to identify neural correlates that are not expected, or that do not fully correspond to the brain parcellation provided by an anatomical atlas.

In order to integrate findings of previous investigations into a bigger, coherent picture, the present study thus aimed to identify possible networks underlying the line bisection task, using machine learning-based, multivariate voxel-wise analysis approaches. First, we used structural lesion data and conducted a multivariate lesion-behaviour mapping analysis to detect areas where focal damage might directly induce LBEs. This approach is powerful in identifying complex configurations of neural correlates such as in brain networks (Mah et al., 2014; Sperber et al., 2019b; Zhang et al., 2014). Second, by quantifying virtual white matter disconnection related to lesion location, i.e. depicting white matter connections in healthy brains running through patients’ lesion areas, we further aimed to investigate remote pathological processes.

## 2. Methods

### 2.1 Patient recruitment

The sample consisted of 172 neurological patients admitted to the Centre of Neurology at Tuebingen University, which were screened for a first ever right-hemisphere stroke. The sample size was determined following recommendations from Sperber et al. (2019), concluding that sample sizes of at least 100–120 subjects are required to optimally model voxel-wise lesion location in SVR-LSM. We excluded patients with diffuse or bilateral brain lesions, patients with tumours, as well as patients in whom MRI or CT scans revealed no obvious lesions. As we were interested in the typical rightward LBE, we excluded nine patients with a leftward deviation in the line bisection task of more than 7.91% from the midline (see below), leaving 163 patients that were included in the present analyses. Table 1 shows demographic and clinical data of all 163 patients. 155 of these patients with valid line bisection testing were included in a previous study addressing the anatomy of spatial neglect evaluated with cancellation tasks (Wiesen et al., 2019). We report how we determined our sample size, all data exclusions, all inclusion/exclusion criteria, whether inclusion/exclusion criteria were established prior to data analysis, all manipulations, and all measures in the study. Subjects gave their informed consent to participate in the study. The study was conducted in accordance with the ethical guidelines of the revised Declaration of Helsinki.

**Table 1:** Demographic and clinical data of all 163 patients. For descriptive information it was determined whether a Line Bisection score was in the pathological range; the cut-off was an ipsilesional deviation of >7.77% (Sperber and Karnath, 2016a). Data are represented either i) as mean, with standard deviationin parentheses and range in brackets, or ii) as percent of the sample. Line bisection is reported as percent of deviation from the true midpoint.To determine if Centre of Cancellation (CoC) scores were in the pathological range, cut-offs were set at >.081 for the Bells Cancellation Task and >.083 for the Letter Cancellation test(Rorden and Karnath, 2010). Further we determined if patients with and without pathological scores differed significantly from each other (Mann-Whitney-U for continuous variables or Chi-Square for categorical variables).

|  | Bisection Bias | No Bias | X<sup>2</sup> / U | sig. |
| --- | --- | --- | --- | --- |
| Age (years) | 60.4 (14.0) [18-81] | 61.6 (13.8) [26-93] | 2774 |  |
| Sex (M/F) | 19/26 | 50/68 | 0 |  |
| Aetiology (Ischemia/Haemorrhage) | 38/7 | 101/17 | 0.034 |  |
| Lesion size (cm<sup>3</sup>) | 80.7 (66.9) [2.9-274.8] | 28.5 (31.4) [0.2-144.9] | 1116 | * |
| Time between lesion and assessment (days) | 4.4 (3.9) [0-22] | 4.0 (3.3) [0-17] | 2401 |  |
| Line Bisection Bias (deviation) | 21.1 (18.0) [8-83.2] | 1.2 (3.2) [-7.5-7.6] | 0 | * |
| Bells (% above Cutoff/CoC) | 62.2 / 0.31 (0.32) [-0.03-0.94] | 28.0 / 0.09 (0.18) [-0.11-0.87] | 1426 | * |
| Letter (% above Cutoff/CoC) | 48.8 / 0.26 (0.30) [-0.02-0.89] | 17.0 / 0.08 (0.20) [-0.06-0.96] | 1247 | * |
| Visual Field Defects (N present) | 27 | 89 | 1.673 |  |

\* < .001

### 2.2 Neuropsychological examination

All patients underwent a neuropsychological examination including the Line Bisection Task (Schenkenberg et al., 1980), Letter Cancellation Task (Weintraub and Mesulam, 1985) and Bells Cancellation Task (Gauthier, Louise Dehaut, Francois Joanette, 1989). The two cancellation tasks were administered and evaluated as previously described (Wiesen et al., 2019) and Centre of Cancellation scores (Rorden and Karnath, 2010) were used for descriptive reports (see Table 1). The mean time between stroke-onset and neuropsychological examination was 4.13 days (SD = 3.43 days). Visual field defects were assessed by confrontation technique.

For the line bisection task, a series of 10 horizontally oriented lines was presented and patients were asked to mark the line’s midpoint with a pencil. Each line was 24 cm long, 0.5 cm wide, and presented separately on a sheet of paper. In order to avoid a systematic bias towards the middle of the sheet, half of the lines were drawn from the left margin of the sheet and half of the lines from the right margin. A displacement to the right side from the midpoint was coded positive and a displacement to the left negative. Line bisection deviation was assessed by averaging the distance between the true midpoint of the lines and the position marked by the patient across all 10 trials. Finally, the LBE was calculated as the percentage of deviation in reference to the maximal possible deviation to the left or right (e.g. −12 cm = −100% maximal to the left; 12 cm = 100% maximal to the right). As we were only interested in the rightward, ipsilesional bias, we excluded nine patients with a pathological contralesional deviation. The cutoff we used for pathological deviation to the left has been determined empirically in 44 healthy controls in a previous investigation (<-7.91%; Sperber and Karnath, 2016a). The non-significant correlation between the LBE and the presence or absence of visual field defects (r = 0.043, n.s) further shows that the LBE is not amplified through visual field defects in the present sample.

### 2.3 Imaging and lesion mapping

Structural imaging was acquired either by MRI (n = 82) or CT (n = 81), performed on average 3.3 days (SD = 4.5 days) after stroke-onset. If both imaging modalities were available, MR scans were preferred over CT scans. Lesions in the individual MR or CT scans were manually marked on transversal slices of the individual scan using MRIcron (www.mccauslandcenter.sc.edu/mricro/mricron). Normalization of CT or MR scans was performed using Clinical Toolbox (Rorden et al., 2012) in SPM8 (www.fil.ion.ucl.ac.uk/spm), which provides age-specific templates in MNI space for both CT and MR scans. If available, MR scans were co-registered with a high resolution T1-weighted structural scan in the normalization process. Fig. S1 shows a descriptive lesion overlap plot of all 163 patients and Fig. 1A of all 163 patients after application of the minimum lesion affection criterion of ∼5%.

**Figure 1:**
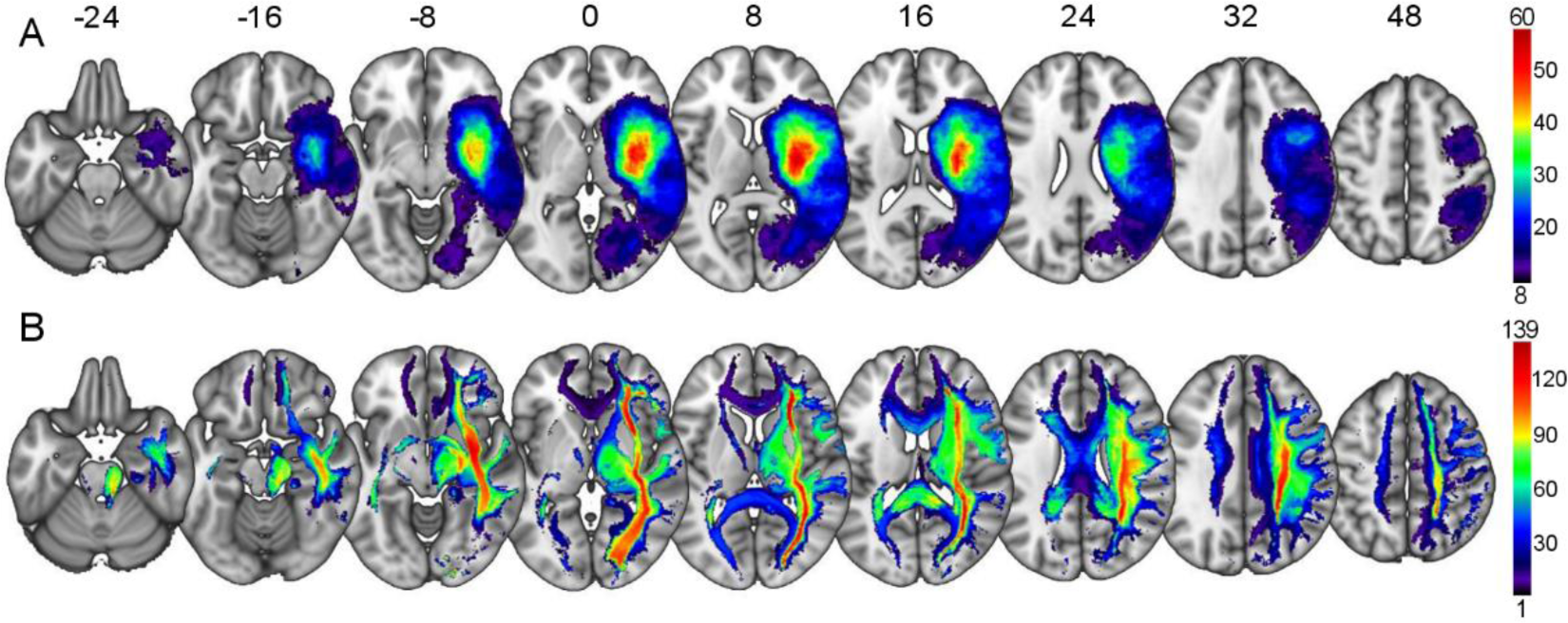
Topography of brain lesions and disconnections. **A**: Lesion overlap plot showing for each voxel the number of patients having a lesion at that location. Only voxels within the voxel mask for statistical testing with at least 8 (∼5%) patients having a lesion are shown. The colourbar indicates the number of overlapping lesions (the peak of N = 60 represents 37% of the total sample). **B**: Lesion disconnection overlap plot of all 163 binarized disconnection probability maps, showing for each voxel the number of patients supposed to have a white matter disconnection at that location. A disconnection is assumed if the probability of disconnection surpasses 50% of being affected in the reference control sample. The colourbar indicates the number of overlapping disconnections (the peak of N = 139 represents 85% of the total sample). Numbers above slices indicate z-coordinates in MNI space. A lesion overlap plot of all patients can be found in the supplementary material (Fig. S1).

For the SVR-LSM analysis, lesion maps were vectorised while applying direct total lesion volume control (dTLVC; see Zhang et al., 2014), which is required in SVR-LSM (Zhang et al., 2014). In the present sample, lesion volume and behaviour showed a significant correlation of 0.50 (p < .001). For further analysis, a matrix with rows representing each case and columns representing the lesion status of each individual voxel was created.

To investigate white matter disconnection, we calculated disconnection maps using the BCBtoolkit (http://toolkit.bcblab.com; Foulon et al., 2018). Each individual lesion mask was used to identify fibre tracks passing through the lesioned area in a set of diffusion weighted imaging datasets of 10 healthy controls (Rojkova et al., 2016), similar to an approach previously used by Kuceyeski et al. (2013). In short, the normalized lesions of each patient were registered to every controls’ native space using affine and diffeomorphic deformations (Avants et al., 2011; Klein et al., 2009). Next, tractography in Trackvis (Wang et al., 2007) was carried out while using the lesion masks as seed. Individual tractographies were then used to create visitation maps, assigning at each voxel a value of 0 or 1 depending on whether the voxel fell into the streamlines of the tract (Thiebaut de Schotten et al., 2011a). These maps were further registered to the original normalized MNI space using the inverse of the precedent deformations. Finally, we calculated for each subject a percentage overlap map by summing at each voxel the normalized visitation map. The resulting disconnection map then indicates for each voxel a probability of disconnection from 0 to 100%, considering the interindividual variability of tract reconstructions in controls (Thiebaut de Schotten et al., 2015). For the subsequent statistical analysis, only voxels with a probability of disconnection of at least 50% were used as input features to predict the pathological behaviour. Fig. 1B shows a descriptive overlap plot of all binarized disconnection maps. We further provide overlap plots for the lesion maps and binarized disconnection maps based on scan modalities in the supplementary material (Fig. S2) showing no substantial differences related to MR or CT lesion delineation. Statistical testing (Mann-Whitney-U) further found no significant differences in lesion volume (*U* = 2112.5, n.s.) between lesions delineated by MR (M = 43.89, SD = 45.69) and lesions delineated by CT (M = 42.16, SD = 53.82).

The procedure for the estimation of the disconnection maps was based on the original lesions as masks for the tractography. It follows that the number of lesioned streamlines should be strongly related to lesion volumes of the seed masks, which can be confirmed in the present sample by the correlation between the number of affected voxels in the disconnection maps (hereafter labelled as ‘disconnection size’) and lesion volumes of the original lesion maps (r = 0.83; p < .001). Further the disconnection size is significantly related to the behavioural outcome (r = 0.39; p <.001). To account for this, each individual disconnection map was read into a vector including dTLVC (Direct total lesion volume control; see Zhang et al., 2014), which corrects for disconnection size in the same way as for the structural lesion map analysis above, ensuring comparability between the two analysis approaches. For the further analysis, a matrix with rows representing cases and columns representing the disconnection status of each individual voxel was used.

### 2.4 Lesion analysis

Our analyses were carried out with a multivariate method that has recently gained popularity in the field of lesion-symptom mapping, namely SVR-LSM (Support Vector Regression based Lesion Symptom Mapping). Support vector regression is a supervised machine learning technique (Cortes and Vapnik, 1995; Drucker et al., 1996) which is able to model the continuous relationship between lesion data and behavioural scores. This method has been employed and validated successfully in previous investigations with real lesion-symptom data (Fama et al., 2017; Griffis et al., 2017; Mirman et al., 2015; Wiesen et al., 2019; Zhang et al., 2014) and synthetic data (Sperber et al., 2019b; Zhang et al., 2014). For a detailed description of this method, we refer to the original study by Zhang et al. (2014). Using this same procedure, but with disconnection maps instead of traditional lesion maps, we additionally performed a disconnection-based multivariate analysis, named hereafter Support Vector Regression based Disconnection-Symptom Mapping (SVR-DSM).

All analyses were performed with MATLAB 2018b and libSVM 3.23 (Chang and Lin, 2013), using SVR with an RBF Kernel. We used a publicly available collection of scripts (https://github.com/yongsheng-zhang/ SVR-LSM) employed in the original study by Zhang et al. (2014) and modified them to make effective use of our computational resources. Main functions of the toolbox were not changed (see http://dx.doi.org/10.17632/2hyhk44zrj.2for public access to the modified analysis scripts). A complete guide on how to perform multivariate lesion-symptom mapping analyses based on support vector regression can be found in Karnath et al. (2019).

First, a parameter optimization procedure via grid search was carried out with lesion maps, as well as with the disconnection maps with 5-fold cross-validations to find the C and γ model parameters with the best trade-off between prediction accuracy and reproducibility, as in a previous investigation (Wiesen et al., 2019). This was done by employing a 5-times 5-fold cross-validation scheme, reflecting model quality when 4/5 of the dataset were used for building the model and 1/5 for testing it afterwards on an ‘unknown’ validation subset. Optimized model parameters C and γ were then used in the final SVR-LSM/SVR-DSM analyses.

The SVR β-parameters were derived for each voxel as described by Zhang and colleagues (2014), representing the strength of the association between each voxel’s lesion or disconnection status and the behavioural score. These β-parameters were tested in a voxel-wise permutation algorithm to assess statistical significance by 10000 permutations, controlled by False Discovery Rate (FDR; Benjamini and Yekutieli, 2001) correction at q = 0.05, and using a cluster threshold of 50mm^3^. For the analysis of structural lesion maps, significant results in cortical and subcortical grey matter regions were labelled with reference to the Automatic Anatomical Labelling atlas (AAL; Tzourio-Mazoyer et al., 2002) distributed with MRIcron (www.mccauslandcenter.sc.edu/mricro/mricron). Topographical results located in white matter were assessed using a tractography-based probabilistic fibre atlas (Thiebaut de Schotten et al., 2011b), extended by the SLF segmentations of a further probabilistic fibre atlas (Rojkova et al., 2016) and thresholded at p >= 0.3 before being overlaid on the statistical topography of both the traditional lesion-symptom analysis and the lesion-symptom disconnection analysis. Note that the exact definition of some of these fibres and their sub-segments – especially the SLF and arcuate fasciculus – differ across the literature. When interpreting our results, we followed the definitions provided by the above atlases.

Furthermore, only clusters with at least 50mm^3^ overlap with a labelled region are reported. Hence, there might be clusters larger than 50mm^3^ in size but without at least 50mm^3^ overlap with a labelled area, thus remaining unassigned.

## 3. Results

### 3.1 Parameter optimization

The parameter optimization routine for the structural lesion maps revealed an optimum C = 10 and γ = 2 which resulted in an average cross-validation prediction accuracy r = 0.25 and Reproducibility = 0.85. For the disconnection maps we achieved a similar model performance as for the lesion maps of prediction accuracy r = 0.30 and Reproducibility = 0.91, by using C = 30 and γ = 9.

Prediction accuracy appeared to be smaller than in previous publications using the technique (Sperber et al., 2019a; Wiesen et al., 2019; Zhang et al., 2014). However, it should be noted that, first, the prediction accuracy reflects prediction performance only after accounting for lesion volume, which is a required procedure for obtaining valid SVR-LSM results (DeMarco and Turkeltaub, 2018; Zhang et al., 2014). This also applies to the disconnection analysis, where larger lesions are more likely to increase disconnection rates. Second, cross-validation prediction accuracy is inherently limited in lesion-behaviour data based only on structural imaging (Sperber, 2020), and prediction accuracy values shown by the SVR-LSM/SVR-DSM technique are thought to only capture variance explainable by lesion-deficit relations, but do not consider further non-topographical variables that might explain additional variance.

### 3.2 SVR-LSM of lesion maps

The SVR-LSM analysis of structural lesion maps, FDR corrected at 0.05 and using lesion volume control by dTLVC, revealed only very few supra-threshold voxels (< 20 connected voxels) and hence no interpretable pattern. The statistical topography can be found online (see data and code availability statement for open access to the files). FDR correction strongly depends on the overall signal in the data, and it might constitute an overly conservative correction method in situations with low signal (Karnath et al., 2018). As seen in the cross-validation, the SVR model’s predictive power was indeed low, and considerably lower than in previous studies using the same methods and similar sample sizes (Sperber et al., 2019a; Wiesen et al., 2019). Accordingly, we post-hoc lowered the threshold to a still conservative cut-off of p <0.001, uncorrected for multiple comparisons, but with a minimum cluster extent threshold of 50mm^3^ to further reduce the number of possible false positives. Thereafter, we found supra-threshold voxels in the inferior parietal lobule – especially within the angular gyrus – and a small cluster in the postcentral gyrus to be associated with LBEs. Moreover, a cluster in the pallidum and extending into the caudate nucleus was associated with LBEs. In white matter, voxels in the anterior commissure, the right arcuate fasciculus and the right superior longitudinal fasciculus (SLF II and SLF III) – SLF II intersecting the right angular gyrus – were associated with line bisection deviation. Note that the cluster overlapped with the arcuate fasciculus as delineated in its entirety by Thiebaut de Schotten et al. (2011), but, following our procedures, we were unable to assign the cluster to one of the three sub-segments of the fibre tract as delineated by the same study. An unassigned cluster larger than 50mm^3^ was detected at the white matter/grey matter border of the inferior temporal gyrus. For an exact overview on relevant clusters of voxels, including effect sizes (i.e. β-weights) and peaks, see table 2 and Fig. 2.

**Table 2:** Significant grey and white matter clusters underlying the LBEin SVR-LSM. Labelling of significant right hemispheric grey and white matter areas found by SVR-LSM uncorrected at 0.001 after 10000 permutations and a cluster extent threshold of 50mm^3^ before being overlaid on the corresponding atlas. Grey matter structures were identified with reference to the Automatic Anatomical Labelling atlas (AAL; Tzourio-Mazoyer et al., 2002). White matter structures were identified using a tractography-based probabilistic fibre atlas (Thiebaut de Schotten et al., 2011b), extended by the SLF segmentations of a further probabilistic fibre atlas (Rojkova et al., 2016) with regions of interest defined at a probability of p >= 0.3. Only clusters with at least 50mm^3^ overlap with a corresponding atlas label are reported.

| GM structure<br>(AAL) | Number of voxels<br>(mm<sup>3</sup>) | $\beta$<br>(Mean/SD) | $\beta$<br>(Peak) |
| --- | --- | --- | --- |
| Angular gyrus | 256 | 2.43/0.27 | 3.10 |
| Pallidum | 71 | 3.79/0.84 | 5.55 |
| Inferior parietal lobe | 63 | 2.89/0.28 | 3.55 |
| Postcentral gyrus | 56 | 2.81/0.21 | 3.40 |
| WM structure<br>(Thiebaut de Schotten et al., 2011b) | Number of voxels<br>(mm<sup>3</sup>) | $\beta$<br>(Mean/SD) | $\beta$<br>(Peak) |
| Arcuate fasciculus | 154 | 2.47/0.26 | 3.19 |
| Anterior commissure | 57 | 3.19/0.98 | 5.09 |
| WM structure<br>(Rojkova et al., 2016) | Number of voxels<br>(mm<sup>3</sup>) | $\beta$<br>(Mean/SD) | $\beta$<br>(Peak) |
| Right SLF II | 460 | 2.69/0.54 | 6.04 |
| Right SLF III | 389 | 2.70/0.57 | 5.87 |

**Figure 2:**
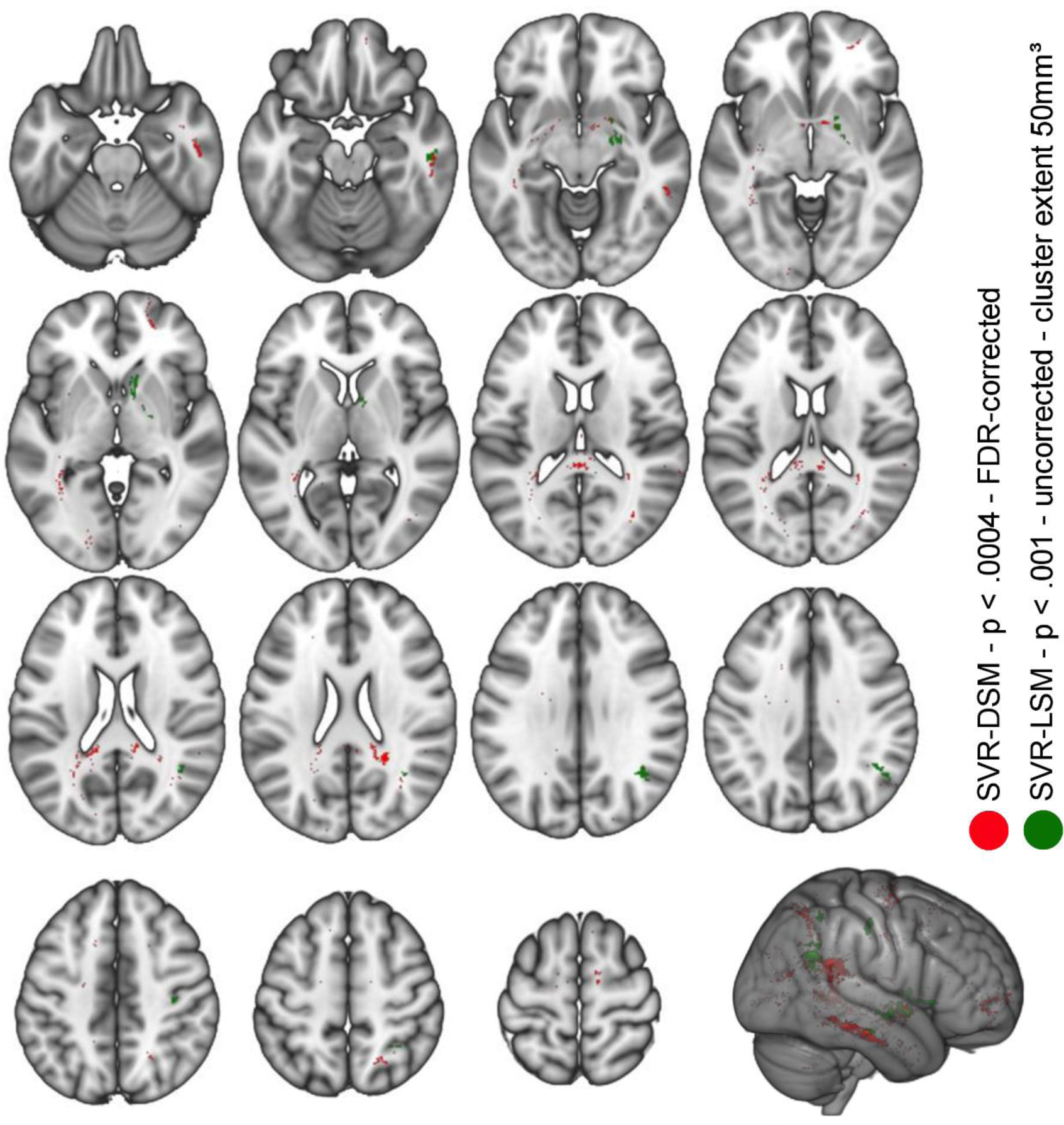
Results of the multivariate lesion-behaviour and disconnection-behaviour mapping. Support vector regression based multivariate lesion-symptom mapping and disconnection-symptom mapping results using data of 163 patients. Green: Permutation-thresholded statistical map of SVR-LSM on line bisection scores (p < 0.001, uncorrected for multiple comparisons), illustrating the anatomical regions significantly associated with the directional LBE. Red: Permutation-thresholded statistical map of SVR-DSM on line bisection scores (p < 0.0004, FDR-corrected at 0.05), illustrating virtual lesion-induced white matter disconnection significantly associated with the directional LBE.

### 3.3 SVR-DSM of disconnection maps

The SVR-DSM analysis of the disconnection maps (Fig. 2), FDR corrected at 0.05 with lesion volume control by dTLVC, showed disconnection to be significantly associated with LBEs in the right hemisphere within the internal capsule and all three branches of the superior longitudinal fasciculus (I, II and III). Moreover, fibres of the posterior corpus callosum and within the anterior commissure were implicated. A further large white matter cluster without corresponding atlas overlap was located close to the lateral part of the right inferior longitudinal fasciculus, between the inferior frontal gyrus and the middle temporal gyrus. For an exact overview on significant areas and effect sizes, see table 3 and Fig. 2.

**Table 3:** Significant white matter clusters underlying the LBE in SVR-DSM. White matter areas where disconnection is associated with the LBE as found by disconnection based SVR-DSM, FDR corrected at 0.05 (p < .0004) based on 10000 permutations. White matter structures were identified using a tractography-based probabilistic fibre atlas (Thiebaut de Schotten et al., 2011b), extended by the SLF segmentations of a further probabilistic fibre atlas (Rojkova et al., 2016) with regions of interest defined at a probability of p >= 0.3. Only clusters with at least 50mm^3^ overlap with a corresponding atlas label are reported.

| WM structure<br>(Thiebaut de Schotten et al., 2011b) | Number of voxels<br>(mm<sup>3</sup>) | $\beta$<br>(Mean/SD) | $\beta$<br>(Peak) |
| --- | --- | --- | --- |
| Corpus Callosum | 209 | 6.48/1.20 | 10 |
| Anterior commissure | 54 | 3.76/1.18 | 5.70 |
| Right internal capsule | 53 | 6.30/2.04 | 9.22 |

| WM structure<br>(Rojkova et al., 2016) | Number of voxels<br>(mm<sup>3</sup>) | $\beta$<br>(Mean/SD) | $\beta$<br>(Peak) |
| --- | --- | --- | --- |
| Right SLF I | 311 | 4.59/1.65 | 9.05 |
| Right SLF II | 263 | 5.25/1.42 | 9.18 |
| Right SLF III | 88 | 3.00/2.12 | 9.18 |

## 4. Discussion

The present study investigated the neural underpinnings of ipsilesional rightward line bisection deviation in acute right hemispheric stroke. We used both multivariate mapping of lesion maps to characterise direct structural damage as well as multivariate mapping of disconnection metrics to reveal additional remote effects of right-hemispheric lesions in both hemispheres.

### 4.1 Grey matter damage related to the line bisection error

By adapting the statistical threshold post-hoc to p <0.001, uncorrected for multiple comparisons, several cortical nodes were found to be involved in line bisection deviation. This included primarily right parietal areas, particularly the inferior parietal lobe, including the angular gyrus, reflecting the importance of posterior brain structures that have been consistently reported by previous studies.

Using lesion overlap plots and subtraction analysis, Binder and colleagues (1992) described LBE as being associated with lesions in the posterior territory of the middle cerebral artery, incorporating the inferior parietal lobe with the angular and supramarginal gyri, as well as posterior parts of the middle temporal gyrus. Following a similar approach, Rorden et al. (2006) were able to replicate the initial findings from Binder et al. (1992) to some extent; they observed the critical area related to LBE at the junction between the middle occipital gyrus and middle temporal gyrus. Kaufmann et al. (2009) used a multiperturbation analysis and reported that the top five areas playing a role for the line bisection task are the supramarginal and angular gyri, the superior parietal lobule, the thalamus and the anterior part of the temporo-parietal junction. Verdon et al. (2010) subsumed a battery of clinical tasks related to spatial attention into different components and found that line bisection mainly loaded on a factor that the authors attributed to perceptual abilities. An anatomical mapping of this component by VLSM implicated a location mainly around the inferior parietal lobe near the supramarginal gyrus. Molenberghs and Sale (2011) were able to replicate this finding by using VLSM and detected a significant cluster related to ipsilesional LBE at the medial part of the right angular gyrus. A slightly different pattern has been shown in the VLSM analysis by Thiebaut de Schotten et al. (2014), who adapted a liberal threshold of p < 0.05 uncorrected for multiple comparisons. The authors not only reported clusters in the superior parietal lobule, the supramarginal gyrus, temporo-parietal junction and the intraparietal sulcus between the angular gyrus and the superior parietal lobe, but also in the middle and inferior frontal gyri as well as the frontal eye fields and the precentral gyrus. Looking at the topographical maps (Thiebaut de Schotten et al., 2014; Fig. 2c), largest effects (i.e. highest z-scores) have nevertheless been detected within the inferior parietal lobule, precentral gyrus, angular and supramarginal gyri, and within the temporo-parietal junction. Interestingly, a pattern including frontal as well as parietal (angular gyrus and superior parietal lobe) and parieto-occipital regions, has been revealed already previously in a VLSM study by Vossel et al. (2011).

The involvement of frontal cortical brain areas in the line bisection task was not confirmed by the present investigation, although our advanced lesion mapping method should be especially suited to find multiple nodes of a network if present. So far only two investigations have reported a direct involvement of frontal cortical areas (Thiebaut de Schotten et al., 2014; Vossel et al., 2011). However, a considerable number of studies described an association of (mostly) caudal parts of fronto-parietal and fronto-occipital white matter pathways (e.g. inferior-longitudinal fasciculus, SLF, arcuate fasciculus and inferior fronto-occipital fasciculus; see below). A possible explanation could be that frontal cortical areas were only rarely affected in some previous studies, and thus not included in the voxel-wise statistical analysis, especially in studies with a small number of cases. Instead, voxels eligible for inclusion into a voxel-wise analysis might have been located in frontal white matter areas, which are more often affected by stroke (cf. Sperber and Karnath, 2016b). Therefore, claims about the absence of frontal involvement in line bisection should be evaluated with caution, as such results might depend on sample characteristics. Accordingly, frontal cortical nodes were only sparsely tested in our analysis due to the exclusion of rarely affected voxels and were mainly restricted to posterior parts of the middle and inferior frontal gyri and the precentral gyrus (see Fig. 1A & Fig. S1).

A recently published multivariate study employing a game theoretical mapping approach showed that the intraparietal sulcus was the main contributor to rightward line bisection deviation (Toba et al., 2017). Additionally, synergistic influences between intraparietal sulcus, temporo-parietal junction and inferior occipital gyrus were reported as being crucial. In a further study, the same authors could delineate areas around the inferior parietal lobe, specifically the angular gyrus and occipital areas, as an anatomical basis of the LBE by VLSM (Toba et al., 2018). Their additional finding of occipital lobe involvement was not confirmed by our investigation.

Besides cortical grey matter influence, our analysis also implicated structural lesions subcortically in the right basal ganglia including the pallidum and extending into the caudate nucleus. The role of damage to the right basal ganglia in spatial attention might be linked to subcortical lesions inducing cortical malperfusion, and, thereby, leading to remote cortical dysfunction (Hillis et al., 2002; Karnath et al., 2005). However, a possible direct effect of subcortical lesions of the basal ganglia to the attentional bias has also been discussed (Parr and Friston, 2018). By simulating attentional deficits within a computational model and affecting the basal ganglia within the model structure, the authors demonstrated direct pathological consequences to the behavioural outcome.

### 4.2 White matter disconnection related to the line bisection error

Our lesion-symptom mapping analysis further showed involvement of the right superior longitudinal fasciculus (SLF II and III) and arcuate fasciculus, while the disconnection-symptom mapping analysis further delineated fibre disruptions within posterior parts (i.e. splenium) of the corpus callosum, and right internal capsule. Further, in the right hemisphere, all three branches of the SLF (SLF I, II & III) overlapped with significant areas of the disconnection topography. An additional non-assigned cluster was located close to the lateral part of the right inferior longitudinal fasciculus at the level of the right inferior temporal gyrus. Several clusters of voxels within the anterior commissure could be delineated by both lesion-symptom mapping and disconnection-symptom mapping.

Besides grey matter areas playing a role for the LBE, damage to white matter connections was reported previously (Golay et al., 2008; Malherbe et al., 2018; Thiebaut de Schotten et al., 2014, 2005; Toba et al., 2018; Vaessen et al., 2016; Verdon et al., 2010). Verdon et al. (2010) described an extension of the significant topography into white matter adjacent to the supramarginal gyrus. Thiebaut de Schotten et al. (2014) showed by track-wise hodological lesion-deficit analysis that the LBE is related to disconnection of the fronto-parietal segment of the arcuate fasciculus and of the second branch of the SLF (II).

When comparing the lesion topography of the VLSM analysis with a common white matter atlas (Thiebaut de Schotten et al., 2011b), Toba and colleagues (2018) reported involvement of the SLF III and the inferior fronto-occipital fasciculus. The authors also evaluated direct fibre tract involvement by using tractography and again found that SLF III integrity predicted rightward line bisection deviation. A descriptive evaluation of the lesion pattern showed that especially caudal disconnections of the SLF III lead to pathological performance. Our study confirmed the latter finding. Initial evidence of the involvement of the SLF comes also from a study using intraoperative electrical stimulation of the SLF and parietal areas during brain surgery (Thiebaut de Schotten et al., 2005). Notably the stimulation of the SLF resulted in greater LBE than stimulation of the right inferior parietal lobe or the posterior superior temporal gyrus, emphasizing again a crucial involvement of fronto-parietal connections in rightward line bisection deviation. With a multivariate approach, Malherbe and colleagues (2018) demonstrated synergetic effects between the SLF and superior temporal gyrus, as well as between the SLF and the inferior fronto-occipital fasciculus contributing to the explanation of leftward line bisection deviation. Vaessen and colleagues (2016) went further and looked directly at DTI-WM metrics. For a factor that loaded on line bisection and text reading, they mapped regions where reduced fractional anisotropy was linked to behavioural deficits. They found two significant clusters that were unaffected by macroscopic lesions, one located in the superior corona radiata adjacent to the trunk of the corpus callosum and the other in the splenium of the corpus callosum. They performed additional fibre tracking, using the clusters above as seeds. In patients scoring worse on this factor, this analysis showed lower track density in parts of the SLF II and III, as well as within the cortico-spinal tract, external capsule and callosal fibres projecting to the left inferior parietal lobe. Taking the second cluster as a seed, again the SLF at the level of the temporo-parietal junction and bilateral projections passing through the splenium of the corpus callosum were reported. Our disconnection analysis showed a similar pattern implicating callosal projection fibres running to the left hemisphere. Correspondingly, a recent intervention study using inhibitory continuous theta burst stimulation (cTBS) over the contralesional parietal lobe found integrity of posterior parts of the corpus callosum to be predictive for successful treatment of directional attentional biases (Nyffeler et al., 2019). Indeed, reduced callosal integrity has been found to be a predictor of persistent attentional deficits in the chronic stage (Lunven et al., 2015), as diagnosed amongst other tests with the line bisection task and leading to persistent symptoms even after therapeutic intervention by prism adaption (Lunven et al., 2019). There is evidence that left parietal areas show an increased BOLD response relative to homologue right hemispheric areas (Corbetta et al., 2005) in right hemisphere stroke patients with attentional deficits. Nyffeler and colleagues (2019) proposed that the pathological hyper-excitability can be normalised after left parietal cTBS and hence, might improve inter-hemispheric communication if callosal fibres are preserved. With respect to the findings of these authors, as well as Lunven and colleagues (2019, 2015), our results indicate that intact callosal fibres might not only be important for neglect recovery and therapy, but that callosal fibre disconnection might also result in an exacerbation of the LBE to the ipsilesional side.

### 4.3 Relation of line bisection and cancellation tasks

It has been repeatedly reported that the directional bisection error dissociates from core symptoms of spatial neglect as measured by different cancellation tasks (Azouvi, 2002; Binder et al., 1992; Ferber and Karnath, 2001; McGlinchey-Berroth et al., 1996; McIntosh et al., 2017; Sperber and Karnath, 2016a; Toba et al., 2017; Verdon et al., 2010). These core symptoms include a spontaneous and sustained deviation of head and eyes to the ipsilesional side and ignoring stimuli on the contralesional side. A simple explanation for the behavioural dissociation between line bisection and cancellation tasks is an anatomical dissociation, i.e. both tasks at least partially rely on different anatomical correlates.

Several previous studies investigated and compared line bisection and cancellation within the same sample. The pioneering work of Binder and colleagues (1992) showed that patients with line bisection errors were likely to have posterior lesions, whereas patients who were impaired on the cancellation task had more anterior damage. This finding has been confirmed by Rorden et al. (2006), and also further studies found dissociations between both tasks (Thiebaut de Schotten et al., 2014; Vaessen et al., 2016; Verdon et al., 2010). Line bisection was associated with lesions to the inferior parietal lobe, posterior parts of the SLF, the arcuate fasciculus and nearby callosal fibres. In contrast, ipsilesional omissions in cancellation tasks were associated with lesions to middle and inferior frontal and superior temporal brain areas, frontal parts of parieto-frontal connections, and fronto-frontal connections (Thiebaut de Schotten et al., 2014; Vaessen et al., 2016; Verdon et al., 2010). On the other hand, Thiebaut de Schotten et al. (2014) also pointed at consistencies between the anamtomy underlying deficits in both tasks. They observed disconnection of fibre tracks to lead to pathological behaviour in both tasks, especially for disruptions of the fronto-parietal segment of the arcuate fasciculus and the SLF II, originating from the angular gyrus and terminating in the middle frontal gyrus (Thiebaut de Schotten et al., 2011a; Wang et al., 2016). In a small sample of 25 patients, Toba and colleagues (2017) delineated for both tasks the intra parietal sulcus as the main contributor of performance and observed synergetic relations between several temporal, parietal, and occipital areas. The same authors also reported differences between both tasks. Only cancellation tasks were additionally related to synergetic interactions between the temporo-parietal junction and inferior frontal gyrus. In a combined structural and diffusion tensor imaging study, Toba et al. (2018) reported a central role of the angular gyrus, the inferior parietal lobe, the third branch of the SLF (SLF III) and the inferior fronto-occipital fasciculus for both behavioural tasks, whereas damage to inferior and middle frontal gyri correlated only with cancellation behaviour. Finally, Malherbe and colleagues (2018) found the inferior parietal lobe being crucial not only in ipsilesional line deviation, but also in contralesional omissions on the bells cancellation task in a left hemispheric patient sample.

Whereas the present study analysed the anatomical contributions to pathological line bisection deviation, the focus in the study by Wiesen and colleagues (2019) was to detect the anatomical correlates of the typical bias of neglect patients in cancellation tasks. Most of the patients (N = 155) in the present work were identical to those who participated already in the study by Wiesen et al. (2019). They found a large cortico-subcortical network, incorporating superior and middle temporal areas, and nearby inferior parietal and occipital structures. Interestingly, results also included frontal cortical areas and adjacent white matter. Albeit the anatomy underlying LBEs appears to differ in general from the anatomy found for the cancellation bias by Wiesen et al. (2019), it is interesting to see that there is also some correspondence between the neural correlates of both deficits. When comparing the present results to the recently published multivariate topography of the neural correlates of spatial neglect (Wiesen et al., 2019), this was especially notable for the parietal involvement and the callosal damage. The latter might indicate that pathological line deviation and the spatial neglect syndrome as measured by cancellation behaviour share some pathophysiological key processes.

### 4.4 Are we measuring the wrong line bisection error?

An explanation for the dissociations between a deficit in line bisection and a deficit in cancellation tasks (Azouvi, 2002; Binder et al., 1992; Ferber and Karnath, 2001; McGlinchey-Berroth et al., 1996; McIntosh et al., 2017; Sperber and Karnath, 2016a; Toba et al., 2017; Thiebaut de Schotten et al., 2014; Vaessen et al., 2016; Verdon et al., 2010) might be the lacking internal validity of the traditional way to administer and score line bisection task. LBEs are traditionally analysed, as in the present study, by measuring the deviation of the mark from the true midpoint. Recent studies challenged this traditional approach and instead proposed an alternative theoretical framework to administer and analyse line bisection (McIntosh et al., 2005, 2017). Within this framework, the locations of the left and right end points of the line and the bisection mark are coded in egocentric positions. Line bisection performance is then measured by assessing the influence of each individual end point location on the position of the mark. This results in two factors, of which one, contrary to traditional line bisection, indeed assesses the core symptom of spatial neglect (McIntosh et al., 2017), as measured also by cancellation tasks. Thus, the traditional line bisection assessment provides a biased and somewhat noisy measure of spatial neglect, which is additionally affected by a second factor potentially related to general attentional capabilities (McIntosh et al., 2017). This partial overlap of deficient cognitive functions in classical line bisection might explain divergent behavioural and anatomical findings in previous studies. Following this theoretical framework, line bisection is a noisy measure of spatial neglect, which explains the surprisingly weak prediction performance found in the present data. According to this interpretation, the mixture of at least two different cognitive functions might hamper the lesion-mapping algorithm to detect the true anatomical key areas related to the directional line bisection deficit. Therefore, it will be crucial to use these new insights of McIntosh et al. (2017) into the line bisection task to derive new study protocols focusing systematically on the anatomical dissociations between these two factors.

### 4.5 Heterogeneity in the anatomical correlates of line bisection errors

Besides methodological differences in the investigation of the line bisection task as a result of using univariate versus multivariate approaches (see introduction section), further factors might have led to differences between the present and previous results, including factors such as line length, line position, or exact task demands (e.g., Doricchi et al., 2005; McIntosh et al., 2005; Cavézian et al., 2012). While it is difficult to explain all these findings from the classical theoretical perspectives on line bisection, the two-component theory of line bisection might do so (McIntosh et al. 2005, 2017). Notably, these different factors likely also underlie some of the variance of anatomical findings in the field. The presentation style of the line bisection task in the present study (ten lines of 24 cm length; five oriented along the right margin of the sheet, five oriented along the left margin of the sheet) likely affected the outcome to some degree, and might have led to differences between the present results and results in previous studies.

Another reason for heterogeneous results on the anatomy of line bisection errors in different studies might be the enigmatic role of visual field defects. A small LBE with a shift of the mark towards the contralesional side – mirroring the LBE typically attributed to deficits in spatial attention – is known to be related to visual field defects alone (Kerkhoff & Bucher, 2008). However, this effect was not found in acute stroke patients (Machner et al., 2009; Sperber & Karnath, 2016a), but instead visual field defects were found to be associated with higher LBEs in acute stroke (Doricchi & Angelelli, 1999; Doricchi et al., 2002; Daini et al., 2002; Sperber & Karnath, 2016a). This effect has been termed an ‘amplification effect’ of visual field defects, while, in fact, no generally accepted theory about the causal relation between both variables currently exists. A simple effect of lesion size, i.e. that larger lesions are both more likely to affect primary vision and more likely to induce high LBEs, is also imaginable. The more pronounced LBE in patients with visual field defects could explain why some studies found damage to more posterior brain regions to underlie LBEs compared to spatial neglect. Patients with primary visual defects typically have damage to posterior brain areas, and at the same time, they suffer from a more severe LBE due to the putative amplification effect. Thus, the statistical anatomo-behavioural signal might be enhanced in these posterior areas.

A further possible candidate to complicate the behavioural measure of LBEs could be contralesional deviation, which is sometimes called ‘ipsilesional neglect’ (Kwon & Heilman, 1991; Kim et al., 1999; Sacchetti et al., 2015; Sperber & Karnath, 2016a). Several studies implicated frontal brain areas for this behavioural finding (Kim et al., 1999; Sacchetti et al., 2015). Contralesional deviation is rather rare compared to the common LBE (Sperber and Karnath, 2016a). In the present study, we excluded cases with pathological contralesional deviation, but it is not known how both behavioural deficits are related and if it is the same cognitive processes that are symmetrically disrupted in both cases. It is unknown if both types of LBEs can interact, and if so, in what way they interact.

## Conclusion and perspective

Our findings underline the importance of a network including several cortical nodes and intra-as well as interhemispheric connections in the emergence of the line bisection error. The use of support vector regression based lesion-symptom disconnection mapping revealed that we might miss relevant structures and connections when we only focus on focal damage. However, according to recent findings by McIntosh et al. (2017), the traditional interpretation of the line bisection task might produce a noisy measure, hindering the statistical algorithms to detect all of the relevant anatomical modules and connections. To resolve further inconsistencies in the literature, future studies should perform lesion-symptom mapping by disentangling spatial and non-spatial attentional components of the line bisection task as pointed out by McIntosh et al. (2017), in order to compare their neural correlates to those resulting from anatomo-behavioural analyses of similar tasks, such as cancellation tasks.

## Acknowledgments

This work was supported by the Deutsche Forschungsgemeinschaft (KA 1258/23-1). Daniel Wiesen was supported by the Luxembourg National Research Fund (FNR/11601161). The authors have no competing interests to declare.

## Data and Code Availability Statement

The datasets generated and analyzed during the current study are not publicly available due to the data protection agreement of the Centre of Neurology at Tübingen University, as approved by the local ethics committee and signed by the participants. We provide the scripts of the main analyses, as well as the statistical topographies and overlap maps, available at http://dx.doi.org/10.17632/2hyhk44zrj.2.

## CRediT authorship contribution statement

**Daniel Wiesen**: Conceptualization, Writing – original draft, Formal analysis, Methodology, Investigation. **Christoph Sperber**: Conceptualization, Writing – original draft, Methodology, Investigation. **Hans-Otto Karnath**: Conceptualization, Writing -review & editing.

## Pre-registration Statement

No part of the study procedures was pre-registered prior to the research being conducted. No part of the study analyses was pre-registered prior to the research being conducted.

## Supplementary Material

**Figure S1:**
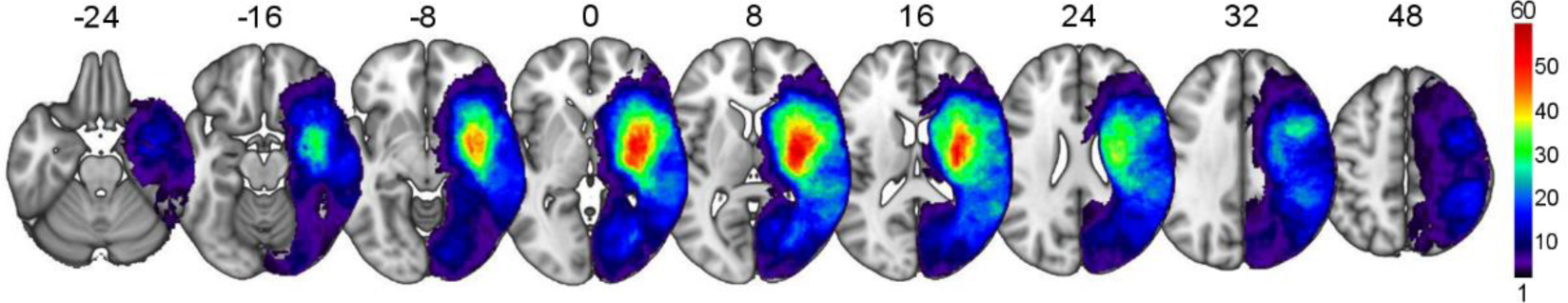
Topography of brain lesions of all patients. Lesion overlap plot of all 163 patients showing for each voxel the number of patients having a lesion at that location. The colour bar indicates the number of overlapping lesions. The peak of N = 60 represents 37% of the total sample. Numbers above slices indicate z-coordinates in MNI space.

**Figure S2:**
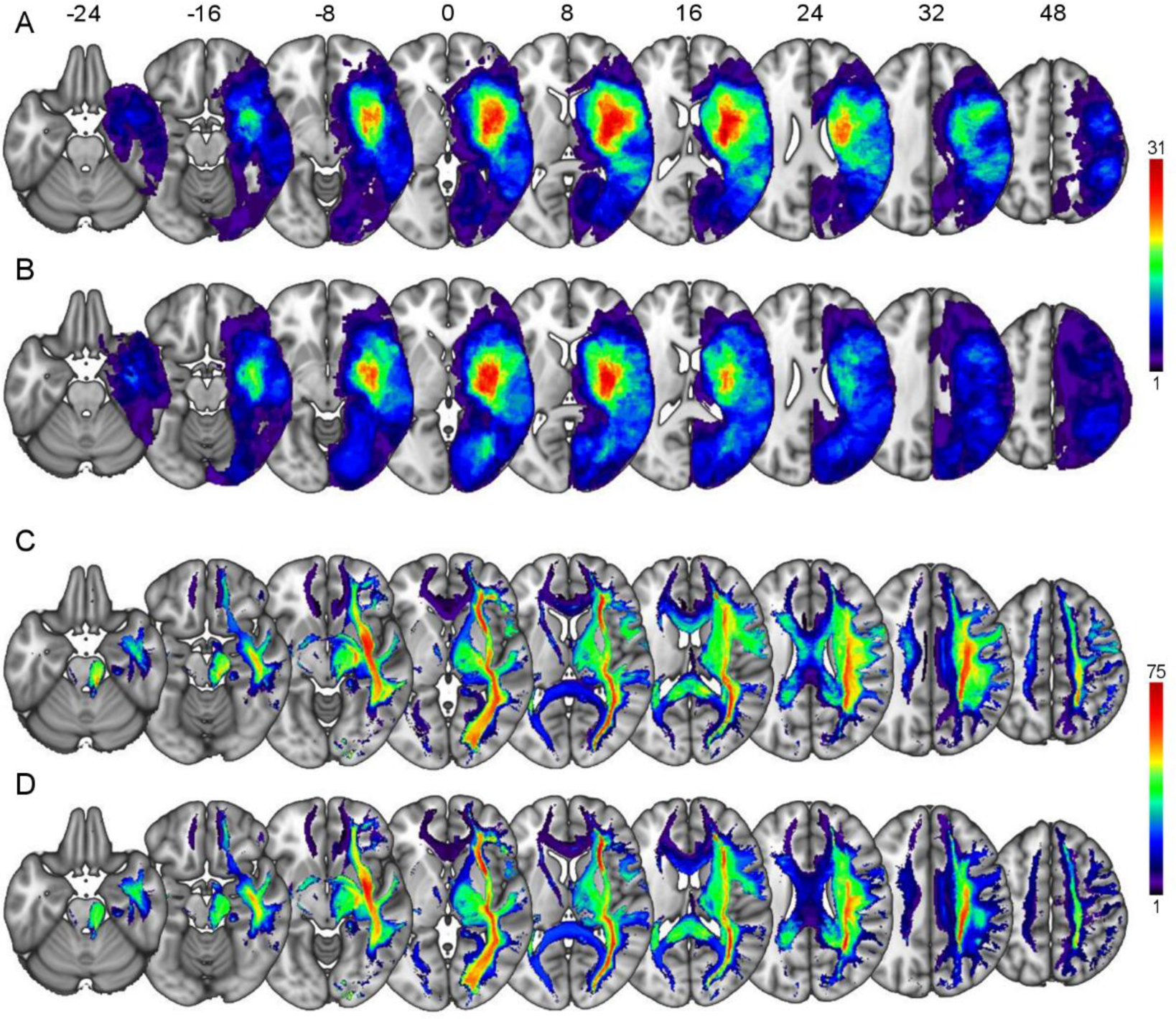
Topography of brain lesions based on scan modality. **A:** Lesion overlap topography of all lesions defined by MR (N = 82). **B:** Lesion overlap topography of all lesions defined by CT (N = 81). **C:** Lesion disconnection overlap plot of all 163 binarized disconnection probability maps, showing for each voxel the number of patients supposed to have a white matter disconnection at that location defined by MR (N = 82). **D:** Lesion disconnection overlap plot of all 163 binarized disconnection probability maps, showing for each voxel the number of patients supposed to have a white matter disconnection at that location defined by CT (N = 81). The colour bar for A and B indicates the number of overlapping lesions. The peak of N = 31 corresponds to ∼37.8% affection of all MR delineated lesions and ∼38.3% affection of all CT delineated lesions. The colour bar for C and D indicates the number of overlapping disconnections. The peak of N = 75 corresponds to 91.5% affection of all MR delineated lesions and 92.6% affection of all CT delineated lesions. Numbers above the slices indicate z-coordinates in MNI space.

## References

Avants, B.B., Tustison, N.J., Song, G., Cook, P.A., Klein, A., Gee, J.C., 2011. A reproducible evaluation of ANTs similarity metric performance in brain image registration. Neuroimage 54, 2033–2044. https://doi.org/10.1016/j.neuroimage.2010.09.025

Azouvi, P., 2002. Sensitivity of clinical and behavioural tests of spatial neglect after right hemisphere stroke. J. Neurol. Neurosurg. Psychiatry 73, 160–166. https://doi.org/10.1136/jnnp.73.2.160

Bartolomeo, P., 2006. A parietofrontal network for spatial awareness in the right hemisphere of the human brain. Arch. Neurol. 63:1238–1241. https://doi.org/10.1001/archneur.63.9.1238

Bartolomeo, P., Thiebaut de Schotten, M., Doricchi, F. 2007. Left unilateral neglect as a disconnection syndrome. Cereb. Cortex 17, 2479–2490.https://doi.org/10.1093/cercor/bhl181

Benjamini, Y., Yekutieli, D., 2001. The control of the false discovery rate in multiple testing under dependency. Ann. Stat. 29, 1165–1188. https://doi.org/10.1214/aos/1013699998

Binder, J., Marshall, R., Lazar, R., Benjamin, J., Mohr, J.P., 1992. Distinct syndromes of hemineglect. Arch. Neurol. 49, 1187–94. https://doi.org/10.1001/archneur.1992.00530350109026

Cavézian, C., Valadao, D., Hurwitz, M., Saoud, M., Danckert, J., 2012. Finding centre: Ocular and fMRI investigations of bisection and landmark taskperformance. Brain Res. 1437, 89–103. https://doi.org/10.1016/j.brainres.2011.12.002

Catani, M., Ffytche, D. H., 2005. The rises and falls of disconnection syndromes. Brain 128, 2224–2239. https://doi.org/10.1093/brain/awh622

Chang, C., Lin, C., 2013. LIBSVM?: A Library for Support Vector Machines. ACM Trans. Intell. Syst. Technol. 2, 1–39. https://doi.org/10.1145/1961189.1961199

Cortes, C., Vapnik, V., 1995. Support-Vector Networks. Mach. Learn. 20, 273–297. https://doi.org/10.1023/A:1022627411411

Daini, R., Angelelli, P., Antonucci, G., Cappa, S.F., Vallar, G., 2002. Exploring the syndrome of spatial unilateral neglect through an illusion of length. Exp. Brain Res. 144, 224–237. https://doi.org/10.1007/s00221-002-1034-8

DeMarco, A.T., Turkeltaub, P.E., 2018. A multivariate lesion symptom mapping toolbox and examination of lesion-volume biases and correction methods in lesion-symptom mapping. Hum. Brain Mapp. 21, 2461–2467. https://doi.org/10.1002/hbm.24289

Doricchi, F., Angelelli, P., 1999. Misrepresentation of horizontal space in left unilateral neglect: Role of hemianopia. Neurology 52, 1845–1852.

Doricchi, F., Galati, G., DeLuca, L., Nico, D., D’Olimpio, F., 2002. Horizontal space misrepresentation in unilateral brain damage. I. Visual and proprioceptive-motor influences in left unilateral neglect. Neuropsychologia 40, 1107–1117.https://doi.org/10.1016/S0028-3932(02)00010-6

Doricchi, F., Guariglia, P., Figliozzi, F., Silvetti, M., Bruno, G., Gasparini, M., 2005. Causes of cross-over in unilateral neglect:between-group comparisons,within-patientdissociations and eye movements. Brain 128, 1386–1406. https://doi.org/10.1093/brain/awh461

Drucker, H., Burges, C.J.C., Kaufman, L., Smola, A., Vapnik, V., 1996. Support vector regression machines. Adv. Neural Inf. Process. Syst. 1, 155–161. https://doi.org/10.1.1.10.4845

Fama, M.E., Hayward, W., Snider, S.F., Friedman, R.B., Turkeltaub, P.E., 2017. Subjective experience of inner speech in aphasia: Preliminary behavioral relationships and neural correlates. Brain Lang. 164, 32–42. https://doi.org/10.1016/j.bandl.2016.09.009

Ferber, S., Karnath, H.O., 2001. How to assess spatial neglect--line bisection or cancellation tasks? J. Clin. Exp. Neuropsychol. 23, 599–607. https://doi.org/10.1076/jcen.23.5.599.1243

Foulon, C., Cerliani, L., Kinkingnéhun, S., Levy, R., Rosso, C., Urbanski, M., Volle, E., de Schotten, M.T., 2018. Advanced lesion symptom mapping analyses and implementation as BCBtoolkit. Gigascience 7, 1–17. https://doi.org/10.1093/gigascience/giy004

Gauthier, L., Dehaut, F., Joanette, Y., 1989. The Bells Test: A quantitative and qualitative test for visual neglect. Int. J. Clin. Neuropsychol. 11, 49–54.

Godefroy, O., Duhamel, A., Leclerc, X., Saint Michel, T., Henon, H., Leys, D., 1998. Brain–behaviour relationships: some models and related statistical procedures for the study of brain-damaged patients. Brain. 121, 1545–56. https://doi.org/10.1093/brain/121.8.1545

Golay, L., Schnider, A., Ptak, R., 2008. Cortical and subcortical anatomy of chronic spatial neglect following vascular damage. Behav. Brain Funct. 4, 1–10. https://doi.org/10.1186/1744-9081-4-43

Griffis, J.C., Nenert, R., Allendorfer, J.B., Szaflarski, J.P., 2017. Damage to white matter bottlenecks contributes to language impairments after left hemispheric stroke. NeuroImage Clin. 14, 552–565. https://doi.org/10.1016/j.nicl.2017.02.019

Halligan, P.W., Cockburn, J., Wilson, B.A., 1991. The behavioural assessment of visual neglect. Neuropsychol. Rehabil. 1, 5–32. https://doi.org/10.1080/09602019108401377

Hillis, A.E., Wityk, R.J., Barker, P.B., Beauchamp, N.J., Gailloud, P., Murphy, K., Cooper, O., Metter, E.J., 2002. Subcortical aphasia and neglect in acute stroke: the role of cortical hypoperfusion. Brain 125, 1094–104. https://doi.org/10.1093/brain/awf113

Karnath, H.-O., 2009. A right perisylvian neural network for human spatial orienting. In: Gazzaniga MS (Ed.) The Cognitive Neurosciences IV. Cambridge, Mass.: MIT Press, 259–268.

Karnath, H.-O., Rorden, C., 2012. The anatomy of spatial neglect. Neuropsychologia 50, 1010–1017. https://doi.org/10.1016/j.neuropsychologia.2011.06.027

Karnath, H.-O., Sperber, C., Rorden, C., 2018. Mapping human brain lesions and their functional consequences. Neuroimage 165, 180–189. https://doi.org/10.1016/j.neuroimage.2017.10.028

Karnath, H.-O., Sperber, C., Wiesen, D., de Haan, B., 2019. Lesion-Behavior Mapping in Cognitive Neuroscience: A Practical Guide to Univariate and Multivariate Approaches, in: Pollmann, S. (Ed.), Spatial Learning and Attention Guidance. Humana Press, New York, pp. 209–238. https://doi.org/10.1007/7657_2019_18

Karnath, H.O., Zopf, R., Johannsen, L., Berger, M.F., Nägele, T., Klose, U., 2005. Normalized perfusion MRI to identify common areas of dysfunction: Patients with basal ganglia neglect. Brain 128, 2462–2469. https://doi.org/10.1093/brain/awh629

Kaufman, A., Serfaty, C., Deouell, L.Y., Ruppin, E., Soroker, N., 2009. Multiperturbation analysis of distributed neural networks: The case of spatial neglect. Hum. Brain Mapp. 30, 3687–3695. https://doi.org/10.1002/hbm.20797

Kenzie, J.M., Girgulis, K.A., Semrau, J.A., Findlater, S.E., Desai, J.A., Dukelow, S.P., 2015. Lesion Sites Associated with Allocentric and Egocentric Visuospatial Neglect in Acute Stroke. Brain Connect. 5, 413–422. https://doi.org/10.1089/brain.2014.0316

Kerkhoff, G., Bucher, L., 2008. Line bisection as an early method to assess homonymous hemianopia. Cortex 44, 200–205. https://doi.org/10.1016/j.cortex.2006.07.00

Kim, M., Na, D.L., Kim, G.M., Adair, J.C., Lee, K.H., Heilman, K.M., 1999. Ipsilesional neglect: behavioural and anatomical features. J. Neurol. Neurosurg. Psychiatry 67, 35–38.http://dx.doi.org/10.1136/jnnp.67.1.35

Klein, A., Andersson, J., Ardekani, B.A., Ashburner, J., Avants, B., Chiang, M.-C., Christensen, G.E., Collins, D.L., Gee, J., Hellier, P., Song, J.H., Jenkinson, M., Lepage, C., Rueckert, D., Thompson, P., Vercauteren, T., Woods, R.P., Mann, J.J., Parsey, R. V., 2009. Evaluation of 14 nonlinear deformation algorithms applied to human brain MRI registration. Neuroimage 46, 786–802. https://doi.org/10.1016/j.neuroimage.2008.12.037

Kuceyeski, A., Maruta, J., Relkin, N., Raj, A., 2013. The Network Modification (NeMo) Tool: Elucidating the Effect of White Matter Integrity Changes on Cortical and Subcortical Structural Connectivity. Brain Connect. 3, 451–463. https://doi.org/10.1089/brain.2013.0147

Kwon, S.E., Heilman, K.M., 1991. Ipsilateral neglect in a patient following a unilateral frontal lesion. Neurology 41, 2001–2004. https://doi.org/10.1212/wnl.41.12.2001

Lunven, M., De Schotten, M.T., Bourlon, C., Duret, C., Migliaccio, R., Rode, G., Bartolomeo, P., 2015. White matter lesional predictors of chronic visual neglect: A longitudinal study. Brain 138, 746–760. https://doi.org/10.1093/brain/awu389

Lunven, M., Rode, G., Bourlon, C., Duret, C., Migliaccio, R., Chevrillon, E., Thiebaut de Schotten, M., Bartolomeo, P., 2019. Anatomical predictors of successful prism adaptation in chronic visual neglect. Cortex 120, 629–641. https://doi.org/10.1016/j.cortex.2018.12.004

Machner, B., Sprenger, A., Hansen, U., Heide, W., Helmchen, C., 2009. Acute hemianopic patients do not show a contralesional deviation in the line bisection task. J. Neurol. 256, 289–290. https://doi.org/10.1007/s00415-009-0148-3

Mah, Y.H., Husain, M., Rees, G., Nachev, P., 2014. Human brain lesion-deficit inference remapped. Brain 137, 2522–2531. https://doi.org/10.1093/brain/awu164

Malherbe, C., Umarova, R.M., Zavaglia, M., Kaller, C.P., Beume, L., Thomalla, G., Weiller, C., Hilgetag, C.C., 2018. Neural correlates of visuospatial bias in patients with left hemisphere stroke: a causal functional contribution analysis based on game theory. Neuropsychologia 115, 142–153. https://doi.org/10.1016/j.neuropsychologia.2017.10.013

McGlinchey-Berroth, R., Bullis, D.P., Milberg, W.P., Verfaellie, M., Alexander, M., D’Esposito, M., 1996. Assessment of neglect reveals dissociable behavioral but not neuroanatomical subtypes. J. Int. Neuropsychol. Soc. 2, 441–451. https://doi.org/10.1017/S1355617700001521

McIntosh, R.D., Ietswaart, M., Milner, A.D., 2017. Weight and see: Line bisection in neglect reliably measures the allocation of attention, but not the perception of length. Neuropsychologia 106, 146–158. https://doi.org/10.1016/j.neuropsychologia.2017.09.014

Mirman, D., Zhang, Y., Wang, Z., Coslett, H.B., Schwartz, M.F., 2015. The ins and outs of meaning: Behavioral and neuroanatomical dissociation of semantically-driven word retrieval and multimodal semantic recognition in aphasia. Neuropsychologia 76, 208–219. https://doi.org/10.1016/j.neuropsychologia.2015.02.014

Molenberghs, P., Sale, M. V., 2011. Testing for Spatial Neglect with Line Bisection and Target Cancellation: Are Both Tasks Really Unrelated? PLoS One 6, e23017. https://doi.org/10.1371/journal.pone.0023017

Molenberghs, P., Sale, M. V., Mattingley, J.B., 2012. Is there a critical lesion site for unilateral spatial neglect? A meta-analysis using activation likelihood estimation. Front. Hum. Neurosci. 6, 1–10. https://doi.org/10.3389/fnhum.2012.00078

Nyffeler, T., Vanbellingen, T., Kaufmann, B.C., Pflugshaupt, T., Bauer, D., Frey, J., Chechlacz, M., Bohlhalter, S., Müri, R.M., Nef, T., Cazzoli, D., 2019. Theta burst stimulation in neglect after stroke: Functional outcome and response variability origins. Brain 142, 992–1008. https://doi.org/10.1093/brain/awz029

Parr, T., Friston, K.J., 2018. The Computational Anatomy of Visual Neglect. Cereb. Cortex 28, 777–790. https://doi.org/10.1093/cercor/bhx316

Rojkova, K., Volle, E., Urbanski, M., Humbert, F., Dell’Acqua, F., Thiebaut de Schotten, M., 2016. Atlasing the frontal lobe connections and their variability due to age and education: a spherical deconvolution tractography study. Brain Struct. Funct. 221, 1751–1766. https://doi.org/10.1007/s00429-015-1001-3

Rorden, C., Bonilha, L., Fridriksson, J., Bender, B., Karnath, H.O., 2012. Age-specific CT and MRI templates for spatial normalization. Neuroimage 61, 957–965. https://doi.org/10.1016/j.neuroimage.2012.03.020

Rorden, C., Fruhmann Berger, M., Karnath, H.-O., 2006. Disturbed line bisection is associated with posterior brain lesions. Brain Res. 1080, 17–25. https://doi.org/10.1016/j.brainres.2004.10.071

Rorden, C., Karnath, H.O., 2010. A simple measure of neglect severity. Neuropsychologia 48, 2758–2763. https://doi.org/10.1016/j.neuropsychologia.2010.04.018

Sacchetti, D.L., Goedert, K.M., Foundas, A.L., Barrett, A.M., 2015. Ipsilesional neglect: behavioral and anatomical correlates. Neuropsychology 29, 183–190.https://doi.org/10.1037/neu0000122

Schenkenberg, T., Bradford, D.C., Ajax, E.T., 1980. Line bisection and unilateral visual neglect in patients with neurologic impairment. Neurology 30, 509–509. https://doi.org/10.1212/WNL.30.5.509

Sperber, C., 2020. Rethinking causality and data complexity in brain lesion-behaviour inference and its implications for lesion-behaviour modelling. Cortex 126, 49–62. https://doi.org/10.1016/j.cortex.2020.01.004

Sperber, C., Karnath, H.O., 2016a. Diagnostic validity of line bisection in the acute phase of stroke. Neuropsychologia 82, 200–204. https://doi.org/10.1016/j.neuropsychologia.2016.01.026

Sperber, C., Karnath, H.O., 2016b. Topography of acute stroke in a sample of 439 right brain damaged patients. NeuroImage Clin. 10, 124–128. https://doi.org/10.1016/j.nicl.2015.11.012

Sperber, C., Wiesen, D., Goldenberg, G., Karnath, H.O., 2019a. A network underlying human higher-order motor control: Insights from machine learning-based lesion-behaviour mapping in apraxia of pantomime. Cortex 121, 308–321. https://doi.org/10.1016/j.cortex.2019.08.023

Sperber, C., Wiesen, D., Karnath, H.O., 2019b. An empirical evaluation of multivariate lesion behaviour mapping using support vector regression. Hum. Brain Mapp. 40, 1381–1390. https://doi.org/10.1002/hbm.24476

Thiebaut de Schotten, M., Dell’Acqua, F., Forkel, S.J., Simmons, A., Vergani, F., Murphy, D.G.M., Catani, M., 2011a. A lateralized brain network for visuospatial attention. Nat. Neurosci. 14, 1245–1246. https://doi.org/10.1038/nn.2905

Thiebaut de Schotten, M., Dell’Acqua, F., Ratiu, P., Leslie, A., Howells, H., Cabanis, E., Iba-Zizen, M.T., Plaisant, O., Simmons, A., Dronkers, N.F., Corkin, S., Catani, M., 2015. From Phineas Gage and Monsieur Leborgne to H.M.: Revisiting Disconnection Syndromes. Cereb. Cortex 25, 4812–4827. https://doi.org/10.1093/cercor/bhv173

Thiebaut de Schotten, M., Ffytche, D.H., Bizzi, A., Dell’Acqua, F., Allin, M., Walshe, M., Murray, R., Williams, S.C., Murphy, D.G.M., Catani, M., 2011b. Atlasing location, asymmetry and inter-subject variability of white matter tracts in the human brain with MR diffusion tractography. Neuroimage 54, 49–59. https://doi.org/10.1016/j.neuroimage.2010.07.055

Thiebaut de Schotten, M., Tomaiuolo, F., Aiello, M., Merola, S., Silvetti, M., Lecce, F., Bartolomeo, P., Doricchi, F., 2014. Damage to white matter pathways in subacute and chronic spatial neglect: a group study and 2 single-case studies with complete virtual “in vivo” tractography dissection. Cereb. Cortex 24, 691–706. https://doi.org/10.1093/cercor/bhs351

Thiebaut de Schotten, M., Urbanski, M., Duffau, H., Volle, E., Lévy, R., Dubois, B., Bartolomeo, P., 2005. Direct evidence for a parietal-frontal pathway subserving spatial awareness in humans. Science 309, 2226–2228. https://doi.org/10.1126/science.1116251

Toba, M.N., Migliaccio, R., Batrancourt, B., Bourlon, C., Duret, C., Pradat-Diehl, P., Dubois, B., Bartolomeo, P., 2018. Common brain networks for distinct deficits in visual neglect. A combined structural and tractography MRI approach. Neuropsychologia 115, 167–178. https://doi.org/10.1016/j.neuropsychologia.2017.10.018

Toba, M.N., Zavaglia, M., Rastelli, F., Valabrégue, R., Pradat-Diehl, P., Valero-Cabré, A., Hilgetag, C.C., 2017. Game theoretical mapping of causal interactions underlying visuo-spatial attention in the human brain based on stroke lesions. Hum. Brain Mapp. 3471, 3454–3471. https://doi.org/10.1002/hbm.23601

Tzourio-Mazoyer, N., Landeau, B., Papathanassiou, D., Crivello, F., Etard, O., Delcroix, N., Mazoyer, B., Joliot, M., 2002. Automated Anatomical Labeling of Activations in SPM Using a Macroscopic Anatomical Parcellation of the MNI MRI Single-Subject Brain. Neuroimage 15, 273–289. https://doi.org/10.1006/nimg.2001.0978

Vaes, N., Lafosse, C., Nys, G., Schevernels, H., Dereymaeker, L., Oostra, K., Hemelsoet, D., Vingerhoets, G., 2015. Capturing peripersonal spatial neglect: An electronic method to quantify visuospatial processes. Behav. Res. Methods 47, 27–44. https://doi.org/10.3758/s13428-014-0448-0

Vaessen, M.J., Saj, A., Lovblad, K.-O., Gschwind, M., Vuilleumier, P., 2016. Structural white-matter connections mediating distinct behavioral components of spatial neglect in right brain-damaged patients. Cortex. 77, 54–68. https://doi.org/10.1016/j.cortex.2015.12.008

Verdon, V., Schwartz, S., Lovblad, K.O., Hauert, C.A., Vuilleumier, P., 2010. Neuroanatomy of hemispatial neglect and its functional components: A study using voxel-based lesion-symptom mapping. Brain 133, 880–894. https://doi.org/10.1093/brain/awp305

Vossel, S., Eschenbeck, P., Weiss, P.H., Weidner, R., Saliger, J., Karbe, H., Fink, G.R., 2011. Visual extinction in relation to visuospatial neglect after right-hemispheric stroke: quantitative assessment and statistical lesion-symptom mapping. J. Neurol. Neurosurg. Psychiatry 82, 862–868. https://doi.org/10.1136/jnnp.2010.224261

Wang, R., Benner, T., Sorensen, A.G., Wedeen, V.J., 2007. Diffusion toolkit: a software package for diffusion imaging data processing and tractography. Proc Intl Soc Mag Reson Med 15, 3720.

Wang, X., Pathak, S., Stefaneanu, L., Yeh, F.-C., Li, S., Fernandez-Miranda, J.C., 2016. Subcomponents and connectivity of the superior longitudinal fasciculus in the human brain. Brain Struct. Funct. 221, 2075–92. https://doi.org/10.1007/s00429-015-1028-5

Weintraub, S., Mesulam, M.-M., 1985. Mental state assessment of young and elderly adults in behavioral neurology, in: Mesulam M-M. (Ed.), Principles of Behavioral Neurology. F.A. Davis Company, Philadelphia, pp. 71–123.

Wiesen, D., Sperber, C., Yourganov, G., Rorden, C., Karnath, H.-O., 2019. Using machine learning-based lesion behavior mapping to identify anatomical networks of cognitive dysfunction: Spatial neglect and attention. Neuroimage 201, 116000. https://doi.org/10.1016/j.neuroimage.2019.07.013

Wiesen, D., 2020. SVR-LSM Code and Results for Wiesen, Karnath & Sperber: Disconnection somewhere down the line: Multivariate lesion-symptommapping of the line bisection error. Mendeley Data, v2http://dx.doi.org/10.17632/2hyhk44zrj.2

Zhang, Y., Kimberg, D.Y., Coslett, H.B., Schwartz, M.F., Wang, Z., 2014. Multivariate lesion-symptom mapping using support vector regression. Hum. Brain Mapp. 35, 5861–5876. https://doi.org/10.1002/hbm.22590

